# Linear models enable powerful differential activity analysis in massively parallel reporter assays

**DOI:** 10.1101/196394

**Authors:** Leslie Myint, Dimitrios G. Avramopoulos, Loyal A. Goff, Kasper D. Hansen

## Abstract

Massively parallel reporter assays (MPRAs) have emerged as a popular means for understanding noncoding variation in a variety of conditions. While a large number of experiments have been described in the literature, analysis typically uses ad-hoc methods. There has been little attention to comparing performance of methods across datasets.

We present the mpralm method which we show is calibrated and powerful, by analyzing its performance on multiple MPRA datasets. We show that it outperforms existing statistical methods for analysis of this data type, in the first comprehensive evaluation of statistical methods on several datasets. We investigate theoretical and real-data properties of barcode summarization methods and show an unappreciated impact of summarization method for some datasets. Finally, we use our model to conduct a power analysis for this assay and show substantial improvements in power by performing up to 6 replicates per condition, whereas sequencing depth has smaller impact; we recommend to always use at least 4 replicates. Together, these results inform recommendations for differential analysis, general group comparisons, and power analysis and will help improve design and analysis of MPRA experiments. An R package is available from the Bioconductor project at https://bioconductor.org/packages/mpra.

## Introduction

Noncoding regions in the human genome represent the overwhelming majority of genomic sequence, but their function remains largely uncharacterized. Better understanding of the functional consequences of these regions has the potential to greatly enrich our understanding of biology. It is well understood that some noncoding regions are regulatory in nature. It has been straightforward to experimentally test the regulatory ability of a given DNA sequence with standard reporter assays, but these assays are low throughout and do not scale to the testing of large numbers of sequences. Massively parallel reporter assays (MPRA) have emerged as a high-throughput means of measuring the ability of sequences to drive expression (White, 2015; Melnikov, Zhang, et al., 2014). These assays build on the traditional reporter assay framework by coupling each putative regulatory sequence with several short DNA tags, or barcodes, that are incorporated into the RNA output. These tags are counted in the RNA reads and the input DNA, and the resulting counts are used to quantify the activity of a given putative regulatory sequence, typically involving the ratio of RNA counts to DNA counts (Figure 1). The applications of MPRA have been diverse, and there have been correspondingly diverse and ad hoc methods used in statistical analysis. There are three broad categories of MPRA applications: characterization studies, saturation mutagenesis, and differential analysis.

**Figure 1.**
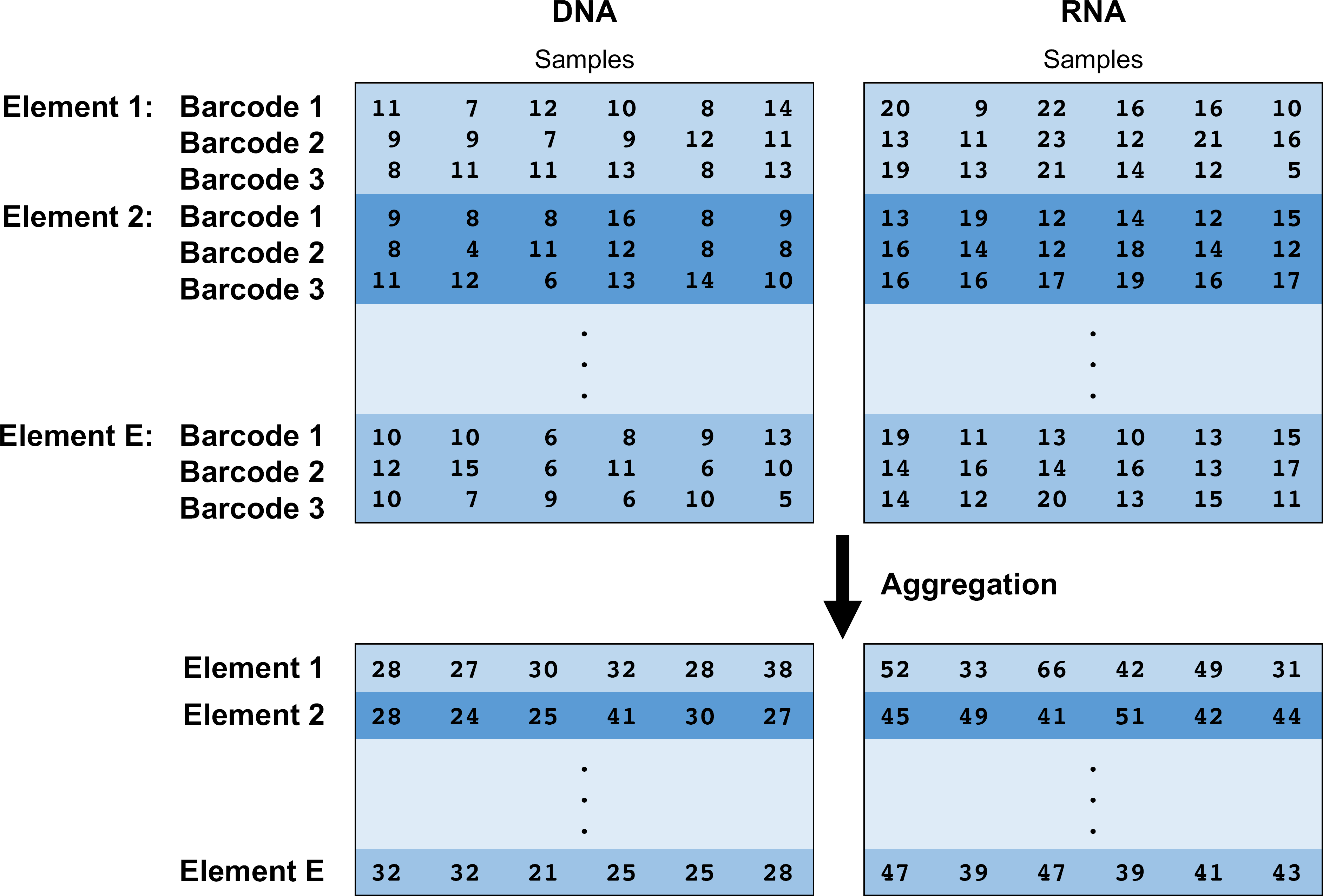
Structure of MPRA data. Thousands of putative regulatory elements can be assayed at a time in an MPRA experiment. Each element is linked to multiple barcodes. A plasmid library containing these barcoded elements is transfected into several cell populations (samples). Cellular DNA and RNA can be isolated and sequenced. The barcodes associated with each putative regulatory element can be counted to obtain relative abundances of each element in DNA and RNA. The process of aggregation sums counts over barcodes for element in each sample. Aggregation is one method for summarizing barcode level data into element level data.

Characterization studies examine thousands of different putative regulatory elements that have a wide variety of sequence features and try to correlate these sequence features with measured activity levels (Grossman et al., 2017; Guo et al., 2017; Safra et al., 2017; Levo et al., 2017; Maricque, Dougherty, and Cohen, 2017; Groff et al., 2016; Ernst et al., 2016; White, Kwasnieski, et al., 2016; Ferreira et al., 2016; Fiore and Cohen, 2016; Farley et al., 2015; Kamps-Hughes et al., 2015; Dickel et al., 2014; Kwasnieski, Fiore, et al., 2014; Mogno, Kwasnieski, and Cohen, 2013; Gisselbrecht et al., 2013; White, Myers, et al., 2013; Smith et al., 2013). Typical statistical analyses use regression to study the impact of multiple features simultaneously. They also compare continuous activity measures or categorized (high/low) activity measures across groups using paired and unpaired t-, rank, Fisher’s exact, and chi-squared tests.

Saturation mutagenesis studies look at only a few established enhancers and examine the impact on activity of every possible mutation at each base as well as interactions between these mutations (Patwardhan, Lee, et al., 2009; Melnikov, Murugan, et al., 2012; Patwardhan, Hiatt, et al., 2012; Kwasnieski, Mogno, et al., 2012; Kheradpour et al., 2013; Birnbaum et al., 2014; Zhao et al., 2014). Analyses have uniformly used linear regression where each position in the enhancer sequence is a predictor.

Differential analysis studies look at thousands of different elements, each of which has two or more versions. Versions can correspond to allelic versions of a sequence (Ulirsch et al., 2016; Tewhey et al., 2016; Vockley et al., 2015) or different environmental contexts (Inoue et al., 2017), such as different cell or tissue types (Shen et al., 2016). These studies have compared different sequence versions using paired t-tests, rank sum tests, and Fisher’s exact test (by pooling counts over biological replicates).

Despite the increasing popularity of this assay, guiding principles for statistical analysis have not been put forth. Researchers use a large variety of ad hoc methods for analysis. For example, there has been considerable diversity in the earlier stages of summarization of information over barcodes. Barcodes are viewed as technical replicates of the regulatory element sequences, and groups have considered numerous methods for summarizing barcode-level information into one activity measure per enhancer. On top of this, a large variety of statistical tests are used to make comparisons.

Recently, a method called QuASAR-MPRA was developed to identify regulatory sequences that have allele-specific activity (Kalita et al., 2017). This method uses a beta-binomial model to model RNA counts as a function of DNA counts, and it provides a means for identifying sequences that show a significant difference in regulatory activity between two alleles. While it provides a framework for two group differential analysis within MPRAs, QuASAR-MPRA is limited in this regard because experiments might have several conditions and involve arbitrary comparisons.

To our knowledge, no method has been developed that provides tools for general purpose differential analysis of activity measures from MPRA. General purpose methods are ones that can flexibly analyze data from a range of study designs. We present mpralm, a method for testing for differential activity in MPRA experiments. Our method uses linear models as opposed to count-based models to identify differential activity. This approach provides desired analytic flexibility for more complicated experimental designs that necessitate more complex models. It also builds on an established method that has a solid theoretical and computational framework (Law et al., 2014). We show that mpralm can be applied to a wide variety of MPRA datasets and has good statistical properties related to type I error control and power. Furthermore, we examine proper techniques for combining information over barcodes and provide guidelines for choosing sample sizes and sequencing depth when considering power. Our method is open https://bioconductor.org/packages/mpra.

## Results

### The structure of MPRA data and experiments

MPRA data consists of measuring the activity of some putative regulatory sequences, henceforth referred to as “elements”. First a plasmid library of oligos is constructed, where each element is coupled with a number of short DNA tags, or barcodes. This plasmid library is then transfected into one or more cellular contexts, either as free-floating plasmids or integrated into the genome (Inoue et al., 2017). Next, RNA output is measured using RNA sequencing, and DNA output as a proxy for element copy number is measured using DNA sequencing (occasionally, element copy number is unmeasured), giving the data structure shown in Figure 1. The log-ratio of RNA to DNA counts is commonly used as an activity outcome measure.

Since each element is measured across a number of barcodes, it is useful to summarize this data into a single activity measure a for a single element in a single sample. Multiple approaches have been proposed for this summarization step. We consider two approaches. First is averaging, where a log-ratio is computed for each barcode, then averaged across barcodes. This treats the different barcodes as technical replicates. The second approach is aggregation, where RNA and DNA counts are each summed across barcodes, followed by formation of a log-ratio. This approach effectively uses the barcodes to simply increase the sequencing counts for that element.

In our investigation of the characteristics of MPRA data we use a number of datasets listed in Table 1. We have divided them into 3 categories. Two categories are focused on differential analysis: one on comparing different alleles and one on comparing the same element in different conditions (retina vs. cortex and episomal vs. chromosomal integration). The two allelic studies naturally involve paired comparisons in that the two elements being compared are always measured together in a single sample (which is replicated). We also use two saturation mutagenesis experiments.

**Table 1.**
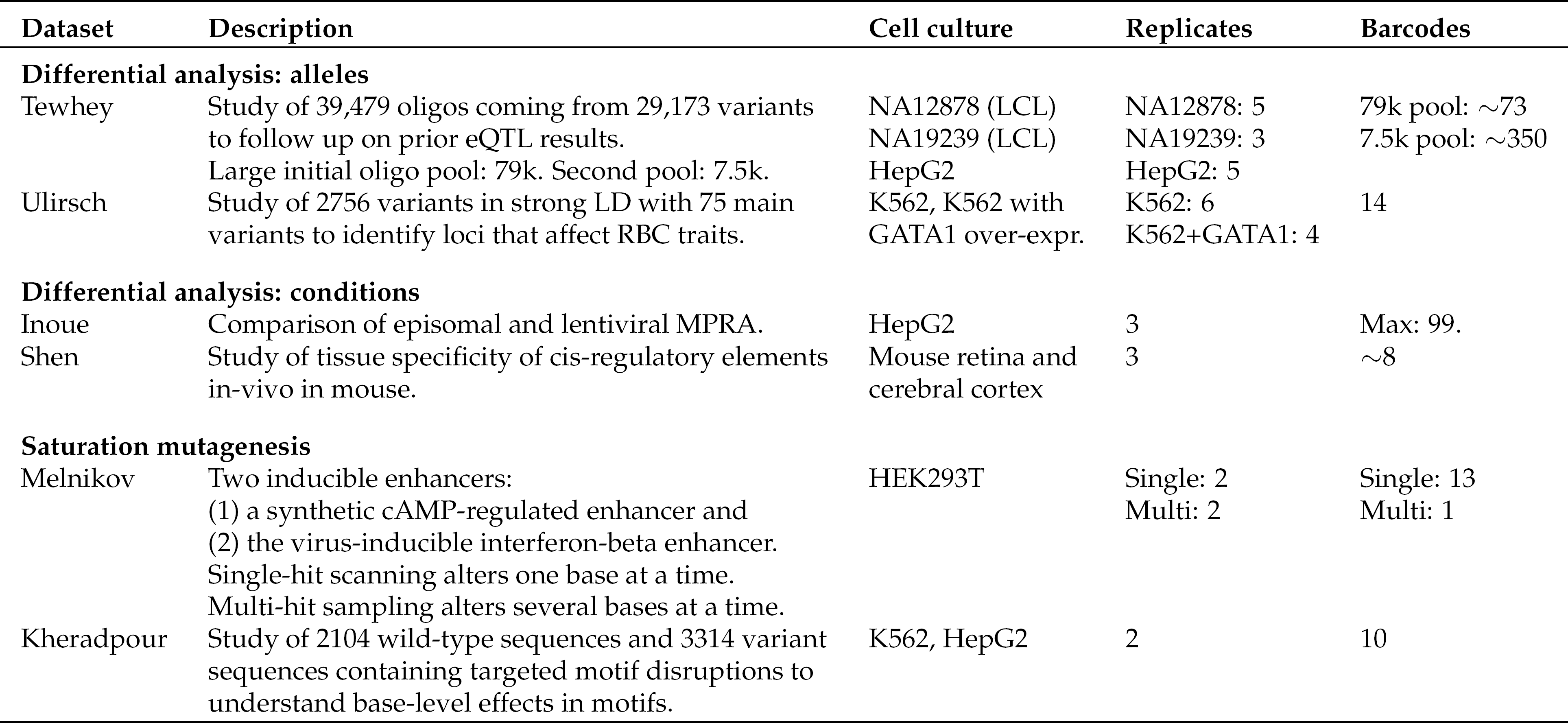
Datasets used for investigations in this paper. All datasets are publicly available.

### The variability of MPRA data depends on element copy number

It is well established that count data from RNA sequencing studies exhibit a mean-variance relationship (McCarthy, Chen, and Smyth, 2012). On the log scale, low counts are more variable across replicates than high counts, at least partly due to inherent Poisson variation in the sequencing process (Marioni et al., 2008; Bullard et al., 2010). This relationship has been leveraged in both count-based analysis methods (Robinson, McCarthy, and Smyth, 2010; Love, Huber, and Anders, 2014) and, more recently, linear model-based methods (Law et al., 2014). For count-based methods, this mean-variance relationship helps improve dispersion estimates, and for linear model-based methods, the relationship allows for estimation of weights reflecting inherent differences in variability for count observations in different samples and genes.

Because MPRAs are fundamentally sequencing assays, it is useful to know whether similar variance relationships hold in these experiments. Due to the construction of MPRA libraries, each element is present in a different (random) copy number, and this copy number ought to impact both background and signal measurements from the element. We are therefore specifically interested in the functional relationship between element copy number and the variability of the activity outcome measure. As outcome measure we use the log-ratio of RNA counts to DNA counts (aggregate estimator), and we use aggregated DNA counts, averaged across samples, as an estimate of DNA copy number. We compute empirical standard deviations of the library size-corrected outcome measure across samples. In Figure 2 we depict this relationship across the previously discussed publicly available datasets (Table 1). For all datasets, with one exception, there is higher variation associated with lower copy number. The functional form is reminiscent of the mean-variance relationship in RNA sequencing data (Law et al., 2014), despite that we here show variance of a log-ratio of sequencing counts.

**Figure 2.**
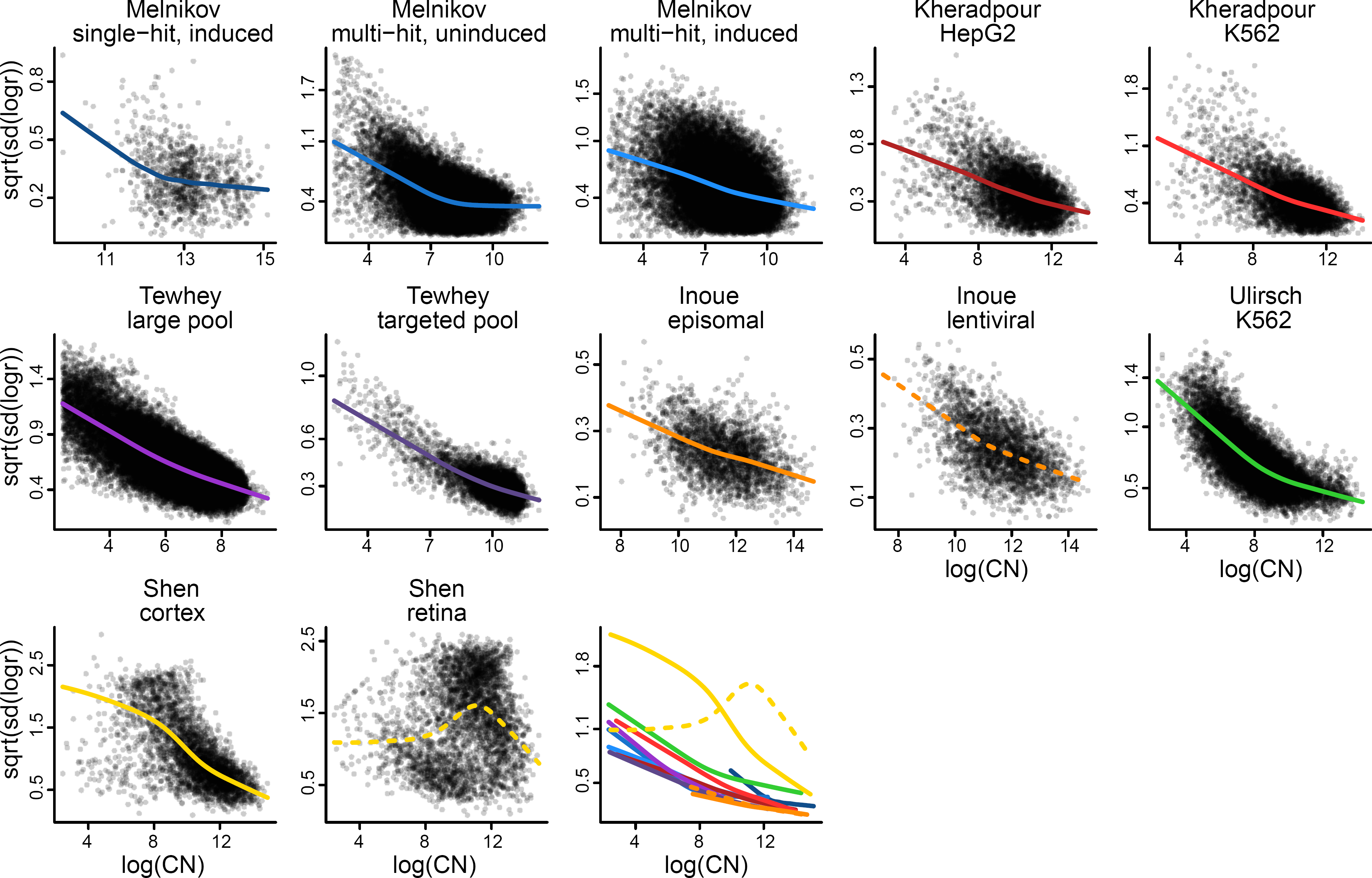
Variability of MPRA activity measures depends on element copy number. For multiple publicly available datasets we compute activity measures of putative regulatory element as the log2 ratio of aggregated RNA counts over aggregated DNA counts. Each panel shows the relationship between variability (across samples) of these activity measures and the average log2 DNA levels (across samples). Smoothed relationships are lowess curves representing the local average variability. The last plot shows all lowess curves on the same figure.

### Statistical modeling of MPRA data

To model MPRA data we propose to use a simple variant of the voom methodology (Law et al., 2014), proposed for analysis of RNA sequencing data. This methodology is based on standard linear models, which are coupled with inverse variance weights representing the mean-variance relationship inherent in RNA sequencing data. The weights are derived from smoothing an empirical mean-variance plot. Similar to voom, we propose to use linear models to model log-ratio activity data from MPRAs, but we estimate weights by smoothing the relationship between empirical variance of the log-ratios and log-DNA copy number, as depicted in Figure 2. This approach has a number of advantages. (1) It is flexible to different functional forms of the variance-copy number relationship. (2) It allows for a unified approach to modeling many different types of MPRA design using the power of design matrices. (3) It allows for borrowing of information across elements using empirical Bayes techniques. (4) It allows for different levels of correlation between elements using random effects. We call this approach mpralm.

The current literature on analysis of MPRA experiments contains many variant methods (see Introduction). To evaluate mpralm, we compare the method to the following variants used in the literature: QuASAR-MPRA, t-tests, and Fisher’s exact test. QuASAR-MPRA is a recently developed method that is targeted for the differential analysis of MPRA data (Kalita et al., 2017). It specifically addresses a two group differential analysis where the two groups are elements with two alleles and uses base-calling error rate in the model formulation. It collapses count information across samples to create three pieces of information for each element: one count for RNA reads for the reference allele, one count for RNA reads for the alternate allele, and one proportion that gives the fraction of DNA reads corresponding to the reference allele. Fisher’s exact test similarly collapses count information across samples. To test for differential activity, a 2-by-G table is formed with RNA and DNA designation forming one dimension and condition designation (with G groups) in the second dimension. The t-test operates on the log ratio outcomes directly; we use the aggregate estimator to summarize over barcodes. Either a paired or unpaired t-test is used based on experimental design.

Both edgeR and DESeq2 are popular methods for analysis of RNA-sequencing data represented as counts. The two methods are both built on negative binomial models, and both attempt to borrow information across genes. These methods allow for the inclusion of an offset. Because both methods use a logarithmic link function, including log-DNA as an offset allows for the modeling of log-ratios of RNA to DNA. This makes these methods readily applicable to the analysis of MPRA data, and they carry many of the same advantages as mpralm. In addition to QuASAR, t-tests, and Fisher’s exact test, we examine the performance of edgeR and DESeq2 for differential activity analysis in our evaluations.

### Simulations shed light on permutation strategies for assessing error rates

Because comparison of type I error rates forms an important part of our methods evaluation (next section), we first present simulation study results regarding the accuracy of permutation procedures for estimating type I error rates. These procedures consist of creating curated null permutations in which the comparison groups are composed of half of the samples from the two original groups. The error rate at different nominal levels is estimated with the median error rate over permutations.

Figure 3 shows how permutation-estimated error rates compare to true type I error rates in a simulation setting with increasing prevalence of differential activity. For all methods we show error estimates resulting from permuting the raw data. For mpralm and the t-test, which operate on the continuous log-ratios, we explore the permutation of residuals proposed in Jiang (2017). We uniformly see that permuting residuals results in substantial overestimation of the error for both methods. Permuting the raw data results in accurate estimation of the error rates in most situations. For this reason, we choose to estimate error rates in real datasets (Table 1) with raw data permutations. We note, however, that permutation of the raw data consistently results in overestimation of QuASAR’s error rates and underestimation of error rates for mpralm for 30% and 50% differential activity. The degree of over- and underestimation increases with the proportion (p) of differential elements, with the effect being more dramatic for QuASAR than for mpralm. We draw on these results when comparing method performance on real datasets in the next section.

**Figure 3.**
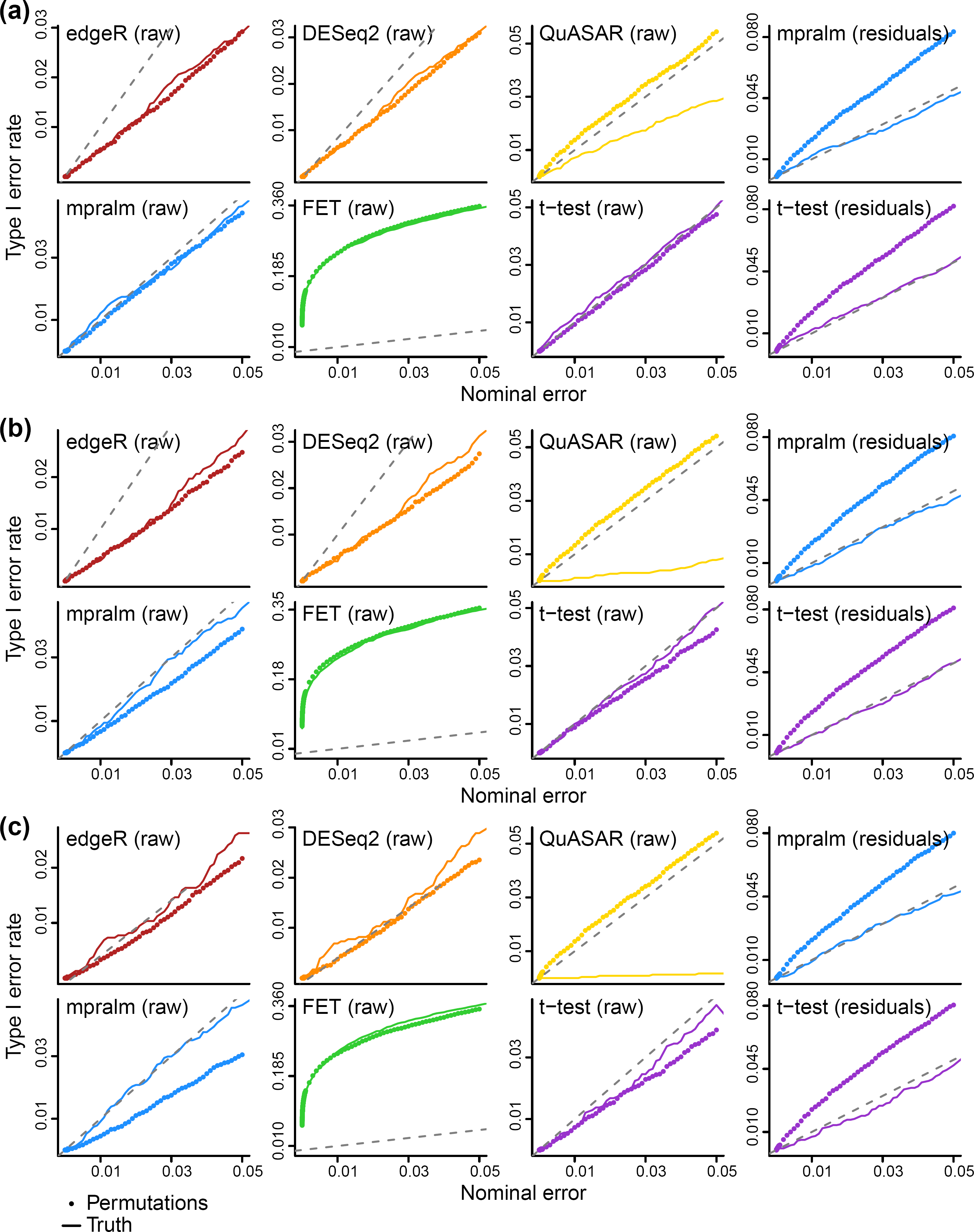
Estimation accuracy of type I error rates using permutations on simulated data. The three sets of panels vary the true proportion (*p*) of elements with differential activity. (a) *p* = 0.1, (b) *p* = 0.3, (c) *p* = 0.5. Each panel shows one method used for differential analysis and compares the true type I error rate to that estimated from null permutations. For mpralm and the t-test, we show error rates from permuting both the raw data and residuals.

### mpralm is a powerful method for differential analysis

First, we focus on evaluating the performance of mpralm for differential analysis. We compare to QuASAR-MPRA, t-tests, Fisher’s exact test, edgeR, and DESeq2. We use four of the previously discussed studies, specifically the Tewhey, Inoue, Ulirsch, and Shen studies. Two of these studies (Tewhey, Ulirsch) focus on comparing the activity of elements with two alleles, whereas the other two (Inoue, Shen) compare the activity of each element in two different conditions. For the allelic studies, we use a random effects model for mpralm and paired t-tests. Both Tewhey et al. (2016) and Ulirsch et al. (2016) compare alleles in different cellular contexts; we observe similar behavior of all evaluations in all contexts (data not shown) and have therefore chosen to depict results from one cellular context for both of these studies. For Tewhey et al. (2016) we depict results both from a large pool of elements used for initial screening and a smaller, targeted pool.

Figure 4 shows p-value distributions that result from running all methods. Across these datasets, all methods except for QuASAR show a well-behaved p-value distribution; high p-values appear uniformly distributed, and there is a peak at low p-values. QuASAR-MPRA consistently shows conservative p-value distributions. We were unable to run QuASAR-MPRA for the Shen dataset. Fisher’s exact test has a very high peak around zero, likely due to the extreme sensitivity of the test with high counts. We examine mpralm using both an average estimator and an aggregation estimator for summarizing across barcodes; this cannot be done for the Tewhey dataset where we do not have access to barcode-level data. To fully interpret these p-value distributions, we need to assess error rates.

To estimate empirical type I error rates, we performed null permutations as described in the previous section. Figure 5 shows estimated error rates (median error rate over the permutations). We observe that Fisher’s exact test has wildly inflated type I error, presumably because the data is overdispersed. QuASAR-MPRA appears well calibrated across datasets, but these error rates might be overestimated. mpralm, t-tests, edgeR, and DESeq2 control the type I error rate but tend to be conservative.

**Figure 4.**
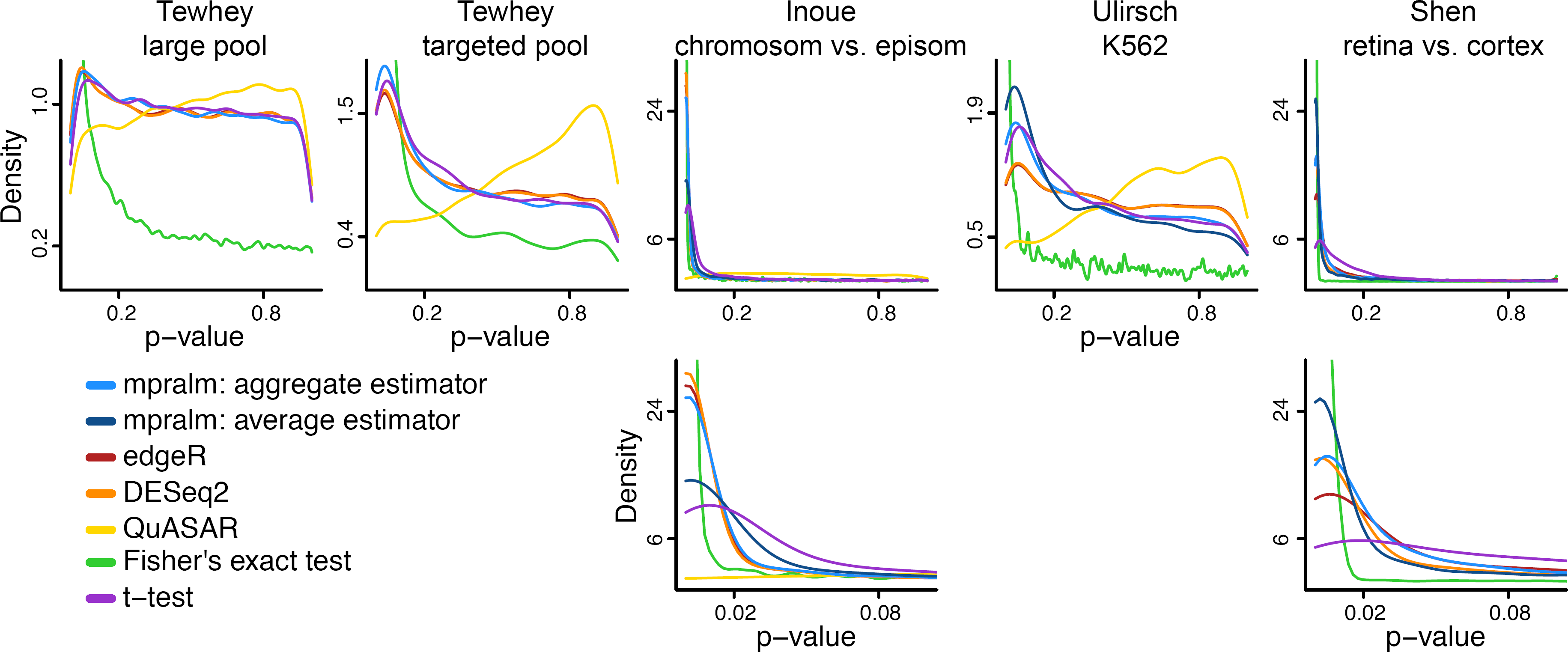
Comparison of detection rates and p-value calibration over datasets. The distribution of p-values for all datasets, including a zoom of the [0,0.1] interval for some datasets. Over all datasets, most methods show p-values that closely follow the classic mixture of uniformly distributed p-values with an enrichment of low p-values for differential elements. For the datasets which had barcode level counts (Inoue, Ulirsch, and Shen), we used two types of estimators of the activity measure (log ratio of RNA/DNA) with mpralm, shown in light and dark blue. We were not able to run QuASAR on the Shen mouse dataset.

**Figure 5.**
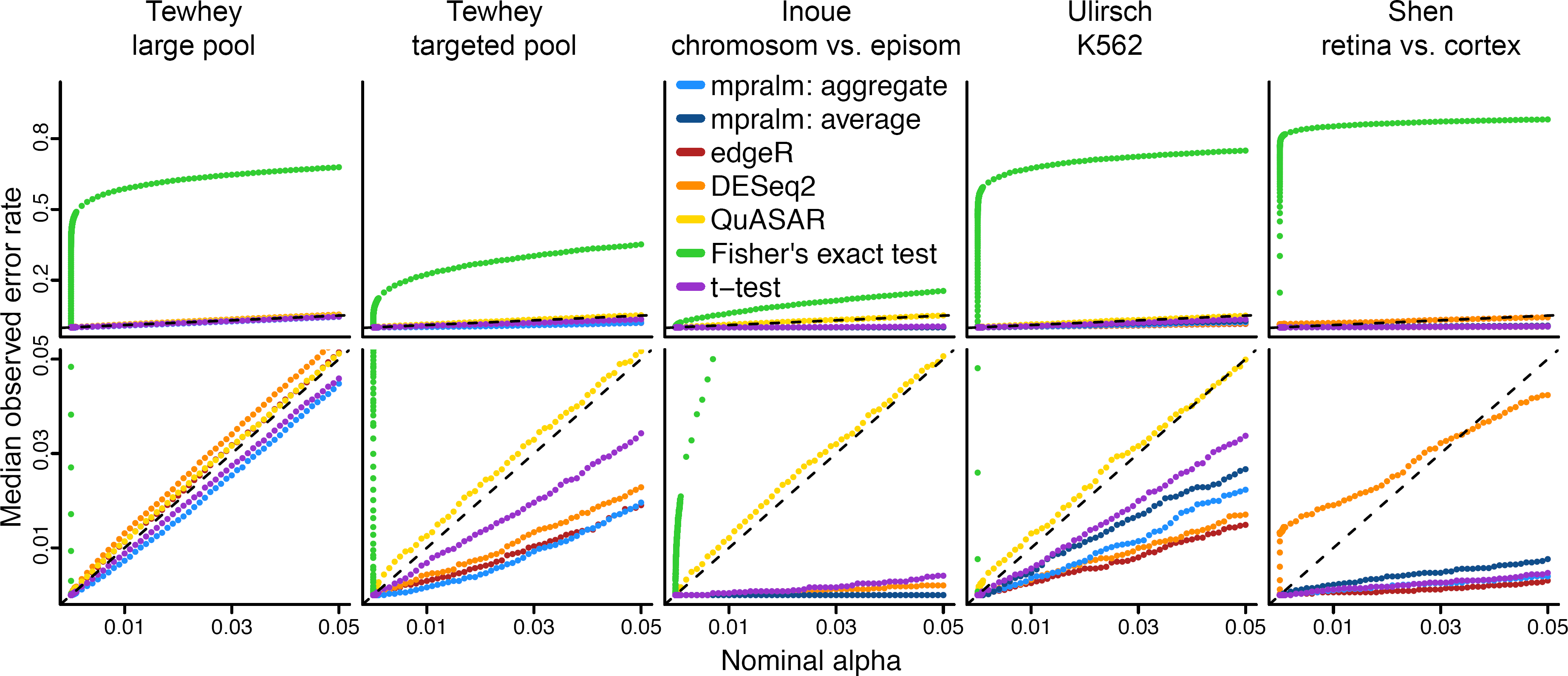
Empirical type I error rates. Type I error rates were estimated for all methods at different nominal levels with null permutation experiments (Methods). For the datasets which had barcode level counts (Inoue, Ulirsch, and Shen), we used two types of estimators of the activity measure (aggregate and average estimator) with mpralm, shown in dark and light blue.

To investigate the trade-off between observed power (number of rejected tests) and type I error rates, we combine these quantities in two ways: (1) we look at the number of rejections as a function of observed type I error rates and (2) we look at estimated FDR as a function of the number of rejections.

In Figure 6 we display the number of rejections as a function of observed type I error rates. In this display, we have essentially used the observed type I error rate displayed in Figure 5 to calibrate the nominal alpha-level. For a fixed error rate, we interpret a high number of rejections to suggest high power. Both Fisher’s exact test and QuASAR-MPRA show poor performance. Because our simulations suggest that the type I error rate of QuASAR can be overestimated with permutations, we expect that it should have better performance than depicted. However, given that its largest number of detections (Figure 6 bottom row) is nearly always as low as the smallest number of detections from other methods, we expect that QuASAR still has poor performance in this regard. Across these datasets, mpralm tends to have the best performance, but edgeR and DESeq2 are competitive. Because our simulations suggest an underestimation of the type I error rate for mpralm, we expect these methods to be closely comparable for this metric.

**Figure 6.**
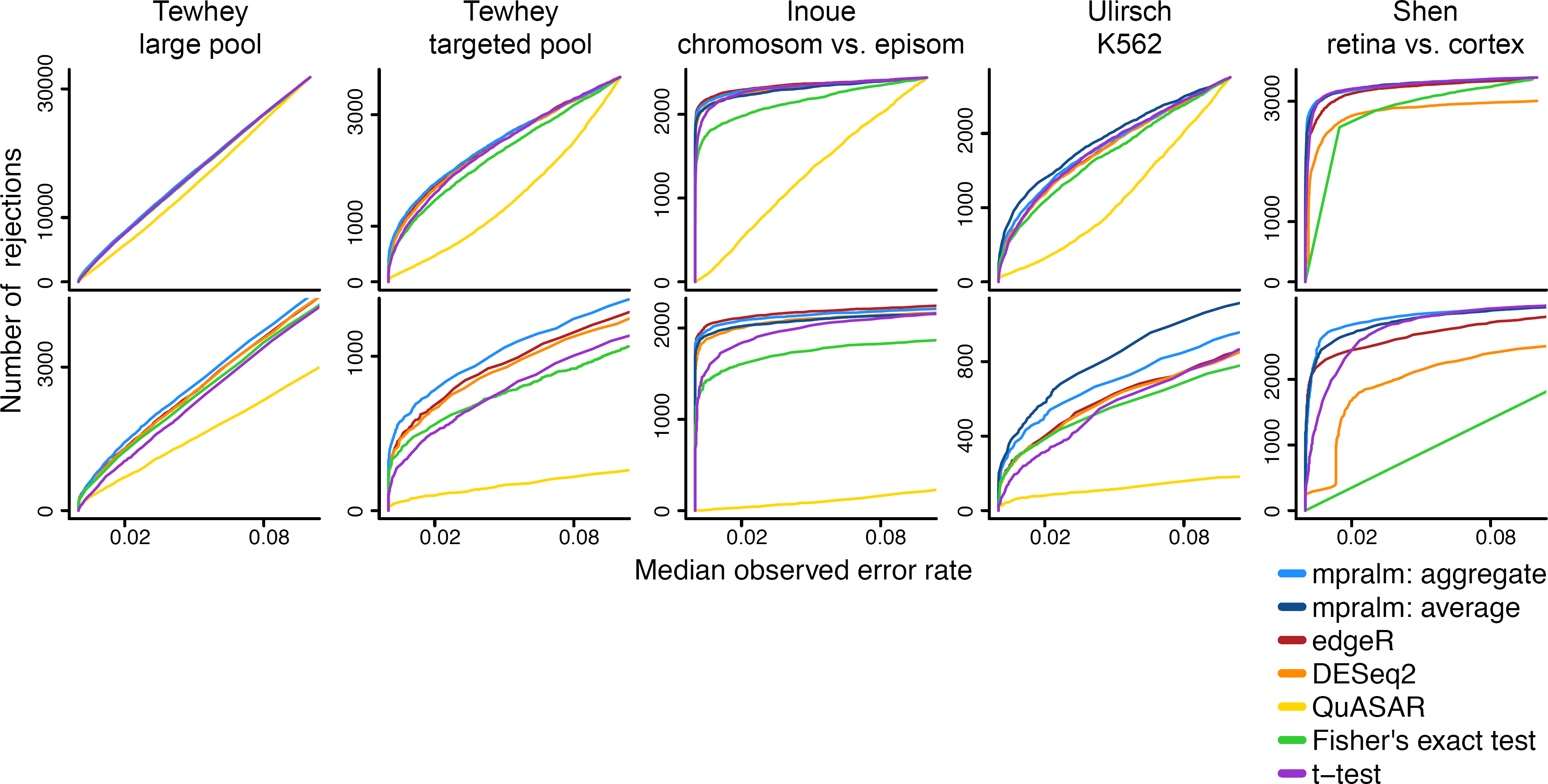
Number of rejections as a function of observed error rate. To compare the detection (rejection) rates of the methods fairly, we compare them at the same observed type I error rates, estimated in Figure 5. The bottom row is a zoomed-in version of the top row. We see that mpralm, edgeR, and DESeq2 consistently have the highest detection rates.

If we know the proportion of true null hypotheses, *π*_0_, we can estimate false discovery rates (FDR). This proportion is an unknown quantity, but we estimate it using a method developed by Phipson (2013) and thereby compute an estimated FDR. In Figure 7 the estimated FDR (for a given *π*_0_) is displayed as a function of the number of rejections. QuASAR-MPRA, t-tests, and Fisher’s exact test tend to have the highest false discovery rates. mpralm tends to have the lowest FDRs. For the Inoue dataset, all methods except for QuASAR have very low FDR, presumably because a very high fraction of elements are expected to be differential given the extreme expected differences between the comparison groups. For this metric, we again expect that QuASAR has better performance than depicted due to error rate overestimation but not enough to be comparable to the other methods. We also expect mpralm to be more comparable to edgeR and DESeq2 given its error rate underestimation.

**Figure 7.**
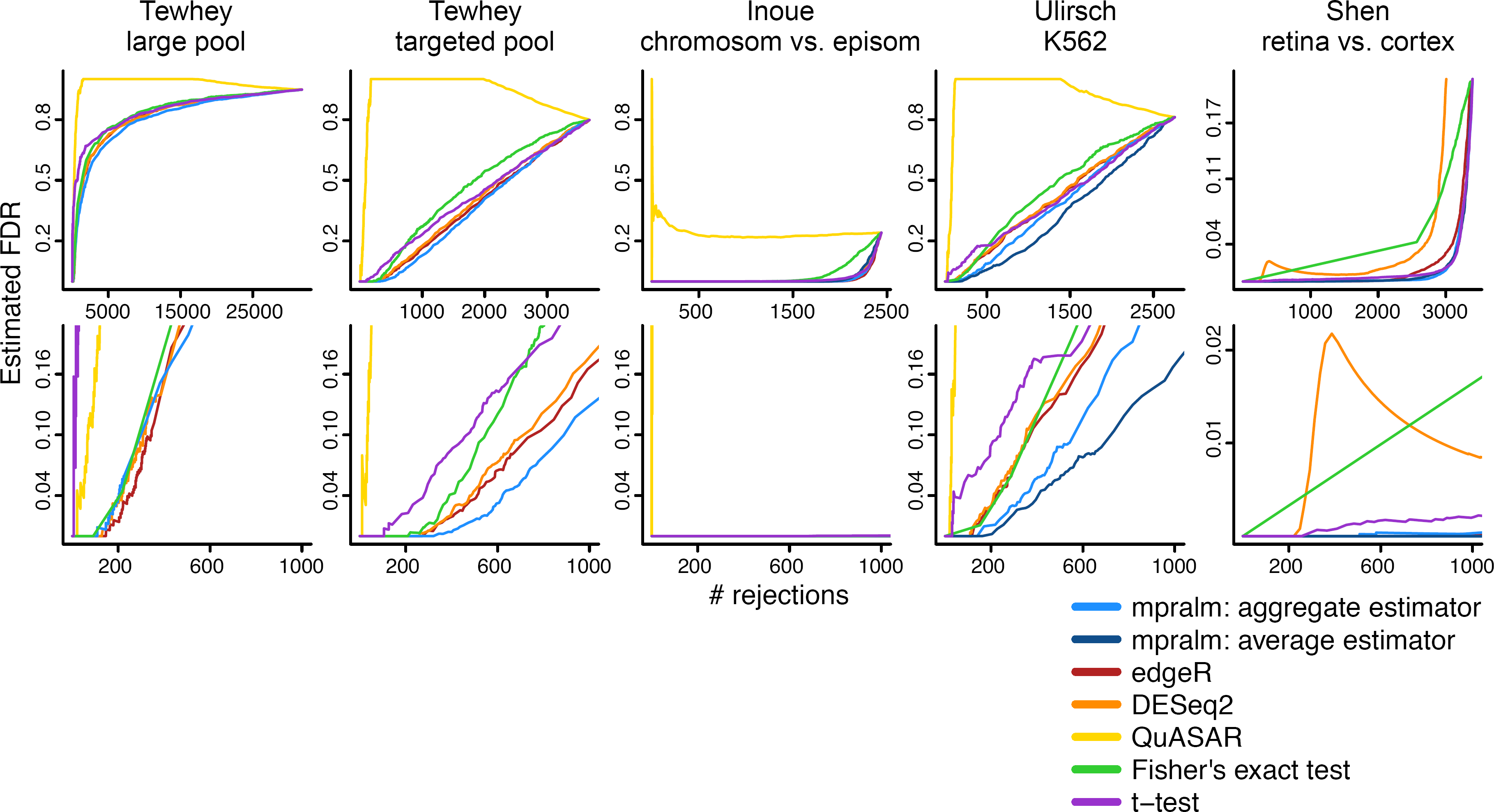
Estimated FDR. For each dataset and method, the false discovery rate is estimated as a function of the number of rejections. This requires estimation of the proportion of true null hypotheses (Supplemental Methods). The bottom row is a zoomed-in version of the top row.

In conclusion, we observe that Fisher’s exact test has too high of an error rate and that QuASAR-MPRA is underpowered; based on these results we cannot recommend either method. T-tests perform better than these two methods but are still outperformed by mpralm, edgeR, and DESeq2, which all have similar performance.

### Comparison of element rankings between methods

While power and error calibration are important evaluation metrics for a differential analysis method, they do not have a direct relation with element rankings, which is often of practical importance. We observe fairly different rankings between mpralm and the t-test and examine drivers of these differences in Figure 8. For each dataset, we find the MPRA elements that appear in the top 200 elements with one method but not the other. We will call these uniquely top ranking elements, and they make up 24% to 64% of the top 200 depending on dataset. For most datasets, DNA, RNA, and log-ratio activity measures are higher in uniquely top ranking mpralm elements (top three rows of Figure 8). It is desirable for top ranking elements to have higher values for all three quantities because higher DNA levels increase confidence in the activity measure estimation, and higher RNA and log-ratio values give a stronger indication that a particular MPRA element has regulatory activity. In the last two rows of Figure 8, we compare effect sizes and variability measures (residual standard deviations). The t-test uniformly shows lower variability but also lower effect sizes for its uniquely top ranking elements. This follows experience from gene-expression studies where standard t-tests tend to underestimate the variance and thereby exhibit t-statistics which are too large, leading to false positives. In MPRA studies, as with most other high-throughput studies, it is typically more useful to have elements with high effect sizes at the top of the list. Such elements are able to picked out in mpralm due to its information sharing and weighting framework.

**Figure 8.**
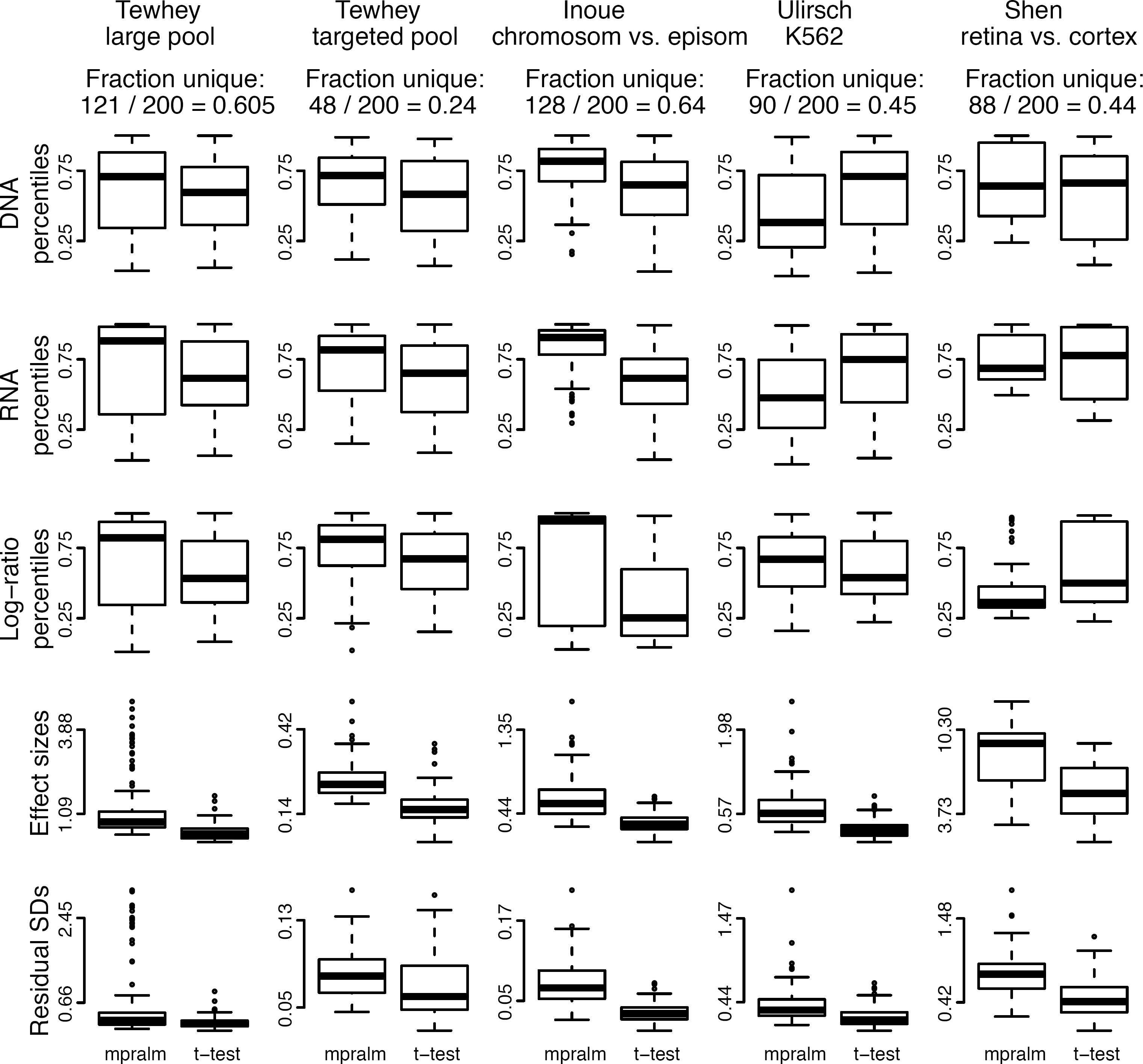
Distribution of quantities related to statistical inference in top ranked elements with mpralm and t-test. MPRA elements that appear in the top 200 elements with one method but not the other are examined here. For these uniquely top ranking elements, the DNA, RNA, and log-ratio percentiles are shown in the first three rows. The effect sizes (difference in mean log-ratios) and residual standard deviations are shown in the last two rows. Overall, uniquely top ranking elements for the t-test tend to have lower log-ratio activity measures, effect sizes, and residual standard deviations.

We similarly compare mpralm rankings with edgeR and DESeq2 rankings in Figures 9 amd 10. The ranking concordance between mpralm and these two methods is much higher than with the t-test. Generally, uniquely top ranking mpralm elements have higher DNA and RNA levels, but lower log-ratio activity measures. Uniquely top ranking mpralm elements also tend to have larger effect sizes. The variability of activity measures (residual SD) is similar among the methods.

**Figure 9.**
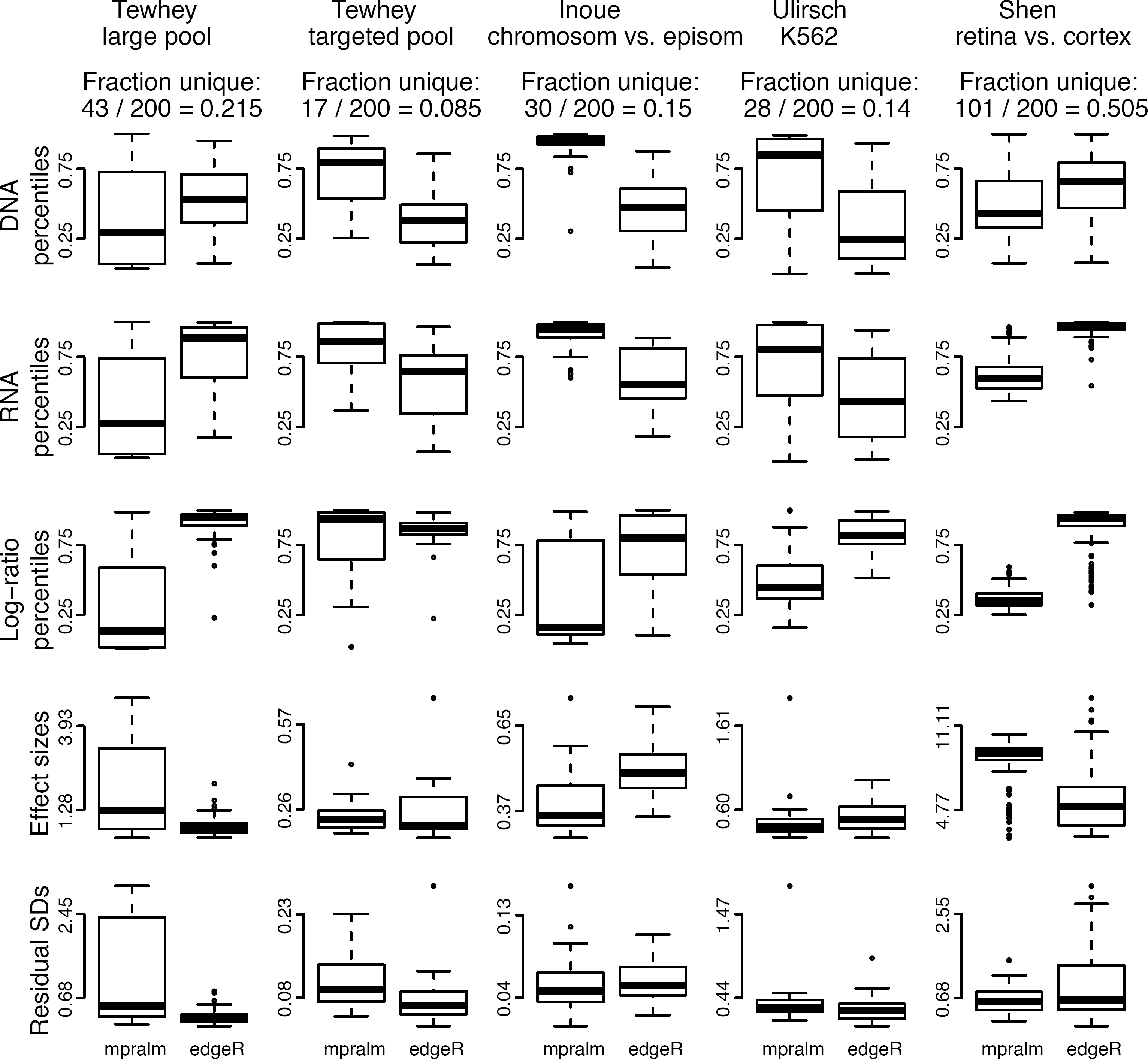
Distribution of quantities related to statistical inference in top ranked elements with mpralm and edgeR. Similar to Figure 8.

**Figure 10.**
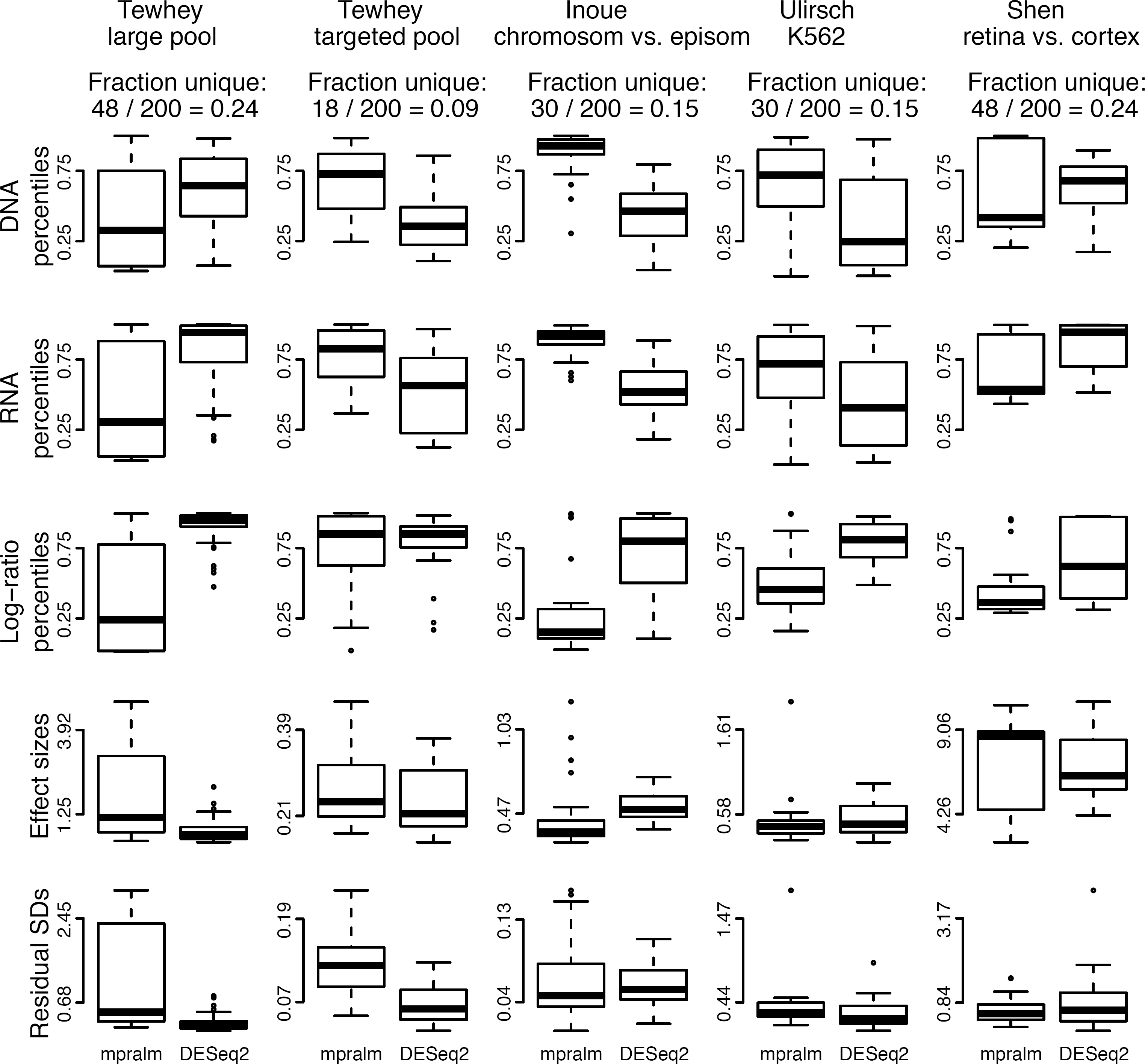
Distribution of quantities related to statistical inference in top ranked elements with mpralm and DESeq2. Similar to Figure 8.

### Accuracy of activity measures and power of differential analysis depends on summarization technique over barcodes

MPRA data initially contain count information at the barcode level, but we typically desire information summarized at the element level for the analysis stage. We examine the theoretical properties of two summarization methods: averaging and aggregation. Under the assumption that DNA and RNA counts follow a count distribution with a mean-variance relationship, we first show that averaging results in activity estimates with more bias. Second, we examine real data performance of these summarization techniques.

Let *R_b_* and *D_b_* denote the RNA and DNA count, respectively, for barcode *b* = 1,…, *B* for a putative regulatory element in a given sample. We suppress the dependency of these counts on sample and element. Typically, *B* is approximately 10 to 15 (for examples, see Table 1). We assume that *R_b_* has mean *μ_r_* and variance *k_r_μ_r_* and that *D_b_* has mean *μ_d_* and variance *k_d_ μ_d_*. Typically the constants *k_d_* and *k_r_* are greater than 1, modeling overdispersion. Negative binomial models are a particular case with *k* = 1 + *ϕμ*, where *ϕ* is an overdispersion parameter. Also let *N_d_* and *N_r_* indicate the library size for DNA and RNA, respectively, in a given sample. Let *p_d_* and *p_r_* indicate the fraction of reads mapping to element *e* for DNA and RNA, respectively, in a given sample so that *μ_r_* = *N_r_p_r_* and *μ_d_* = *N_d_p_d_*. Let *a* be the true activity measure for element *e* defined as *a* := log(*p_r_/p_d_*). When performing total count normalization, the RNA and DNA counts are typically scaled to a common library size *L*.

The average estimator of *a* is an average of barcode-specific log activity measures:

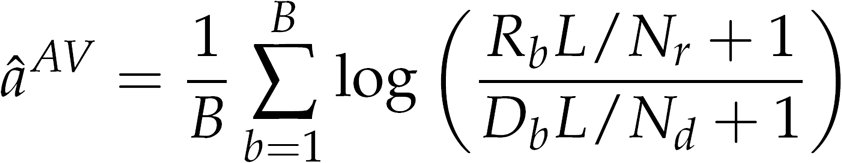

Using a second order Taylor expansion (Supplemental Methods), it can be shown that this estimator has bias approximately equal to

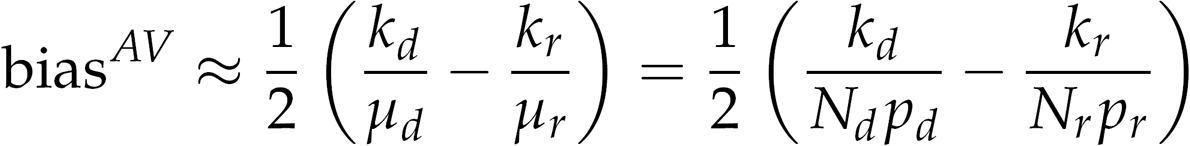

The aggregate estimator of *a* first aggregates counts over barcodes:

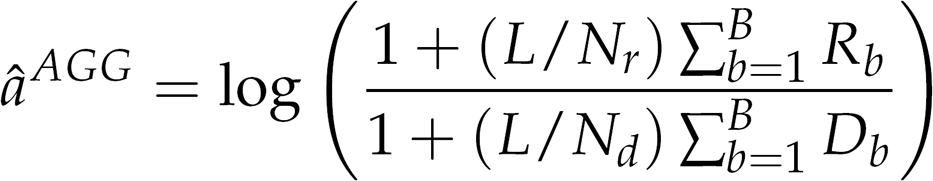

Using an analogous Taylor series argument, we can show that this estimator has bias approximately equal to

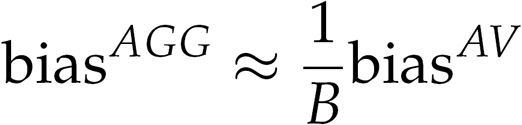

The aggregate estimator has considerably less bias than the average estimator for most MPRA experiments because most experiments use at least 10 barcodes per element. Bias magnitude depends on count levels and the true activity measure *a*. Further, the direction of bias depends on the relative variability of RNA and DNA counts. Similar Taylor series arguments show that the variance of the two estimators is approximately the same.

The choice of estimator can impact the estimated log fold-changes (changes in activity) in a differential analysis. In Figure 11 we compare the log fold-changes inferred using the two different estimators. For the Inoue dataset, these effect sizes are very similar, but there are larger differences for the Ulirsch and Shen datasets.

**Figure 11.**
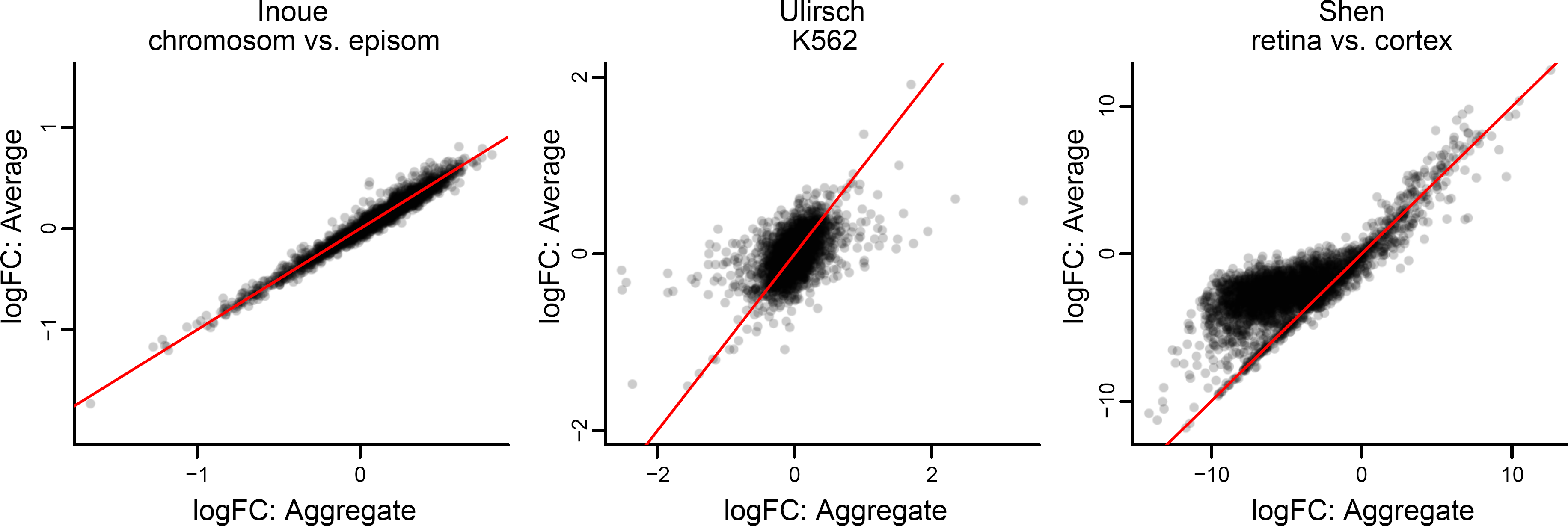
Comparison of the average and aggregate estimators. For the three datasets containing barcode-level information, we compare the effect sizes (log fold changes in activity levels) resulting from use of the aggregate and average estimators. The *y* = *x* line is shown in red.

Aggregation technique affects power in a differential analysis. In the last three columns of Figures 4, 5, 6, and 7, we compare aggregation to averaging using mpralm. The two estimators have similar type I error rates but very different detection rates between datasets. The average estimator is more favorable for the Ulirsch and Shen datasets, and the aggregate estimator is more favorable in the Inoue dataset.

### Recommendations for sequencing depth and sample size

To aid in the design of future MPRA experiments, we used the above mathematical model to inform power calculations. Power curves are displayed in Figure 12. We observe that the variance of the aggregate estimator depends minimally on the true unknown activity measure but is greatly impacted by sequencing depth. We fix one of the two true activity measures to be 0.8 as this is common in many datasets. We use a nominal type I error rate of 0.05 that has been Bonferroni adjusted for 5000 tests to obtain conservative power estimates. We also use ten barcodes per element as this is typical of many studies.

**Figure 12.**
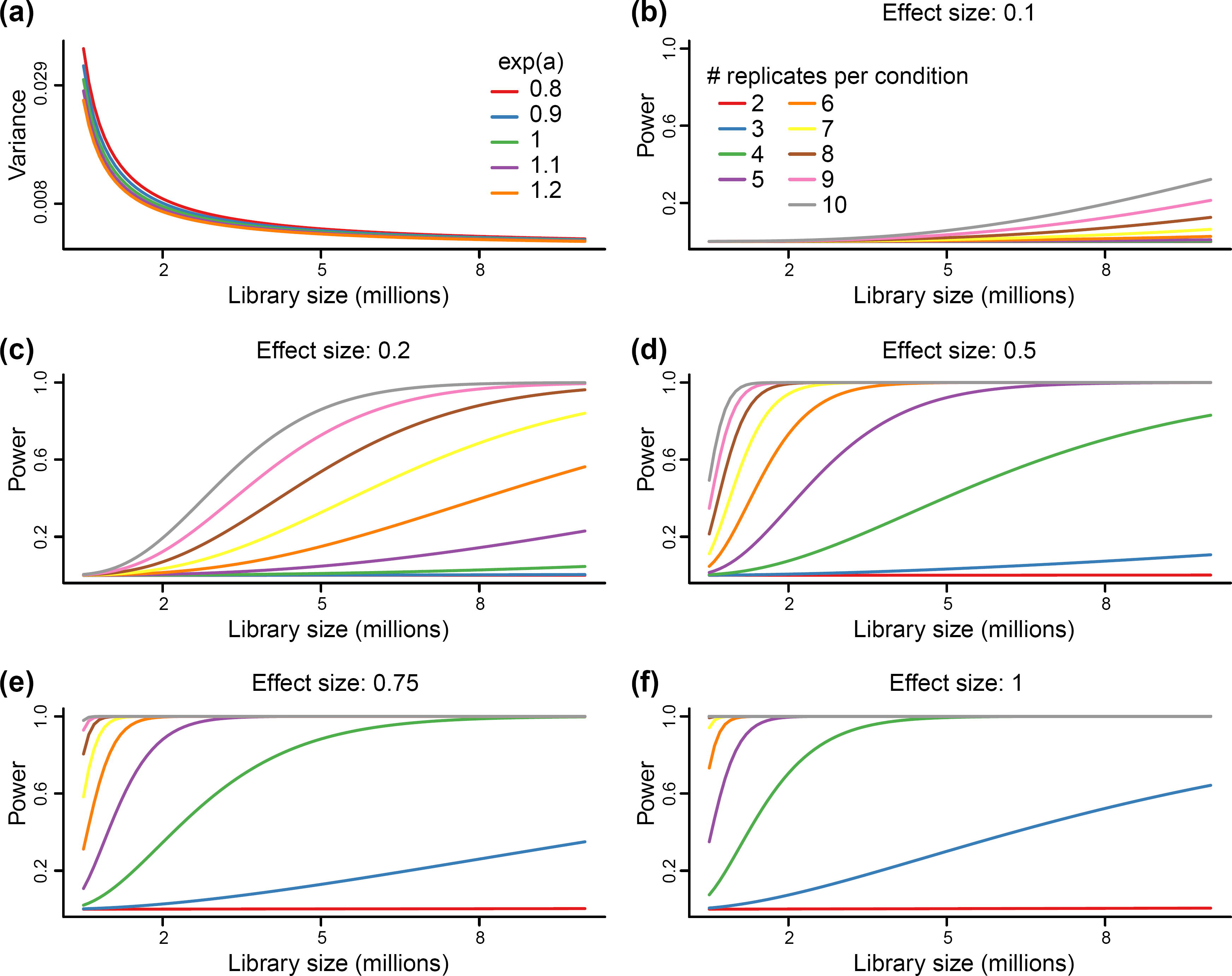
Power analysis. Variance and power calculated based on our theoretical model. **(a)** Variance of the aggregate estimator depends on library size and the true unknown activity level but not considerably on the latter. **(b)**-**(f)** Power curves as a function of library size for different effect sizes and sample sizes. Effect sizes are log_2_ fold-changes.

Our model suggests different impacts of sample size, and a marked impact of increasing the number of replicates, especially between 2 and 6 samples. From Figure 13, we can see that large effect sizes (effect sizes of 1 or greater) are typical for top ranking elements in many MPRA studies. In this situation it is advisable to do 4 or more replicates per group.

**Figure 13.**
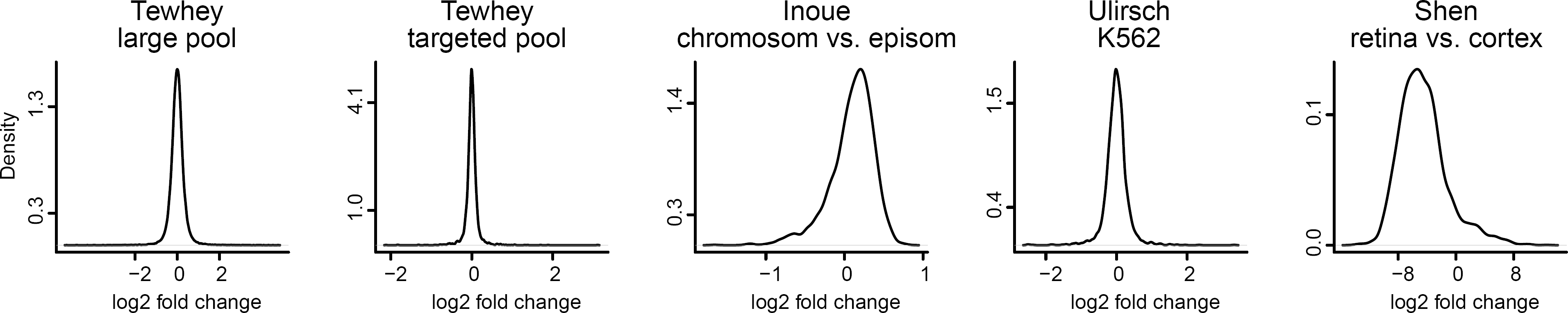
Effect size distributions across datasets. Effect sizes in MPRA differential analysis are the (precision-weighted) differences in activity scores between groups, also called log_2_ fold-changes. The distribution of log_2_ fold changes resulting from using mpralm with the aggregate estimator are shown here.

## Discussion

The field of MPRA data analysis has been fragmented and consists of a large collection of study-specific ad hoc methods. Our objective in this work has been to provide a unified framework for the analysis of MPRA data. Our contributions can be divided into three areas. First, we have investigated techniques for summarizing information over barcodes. In the literature, these choices have always been made without justification and have varied considerably between studies. Second, we have developed a linear model frame-work, mpralm, for powerful and flexible differential analysis. To our knowledge, this is the second manuscript evaluating for statistical analysis in MPRA studies. The first proposed the QuASAR-MPRA method (Kalita et al., 2017), which we show to have worse performance than mpralm. In our comparisons, we provide the largest and most comprehensive comparison of analysis methods so far; earlier work used only two datasets for comparisons. Third, we have analyzed the impact of sequencing depth and number of replicates on power. To our knowledge, this is the first mathematically-based power investigation, and we expect this information to be useful in the design of MPRA studies.

The activity of a regulatory element can be quantified with the log ratio of RNA counts to DNA counts. In the literature, groups have generally taken two approaches to summarizing barcode information to obtain one such activity measure per element per sample. One approach is to add RNA and DNA counts from all barcodes to effectively increase sequencing depth for an element. This is termed the aggregate estimator. Another approach is to compute the log ratio measure for each barcode and use an average of these measures as the activity score for an element. This is termed the average estimator, and we have shown that it is more biased than the aggregate estimator. Because of this bias, we caution against the use of the average estimator when comparing activity scores in enhancer groups (often defined by sequence features). However, it is unclear which of the two estimators is more appropriate for differential analysis.

In addition to barcode summarization recommendations, we have proposed a linear model framework, mpralm, for the differential analysis of MPRA data. Our evaluations show that it produces calibrated p-values and is as or more powerful than existing methods being used in the literature. Its type I error rates appear conservative, so in practice, we recommend performing permutations to estimate error rates.

While the count-based tools, edgeR and DESeq2, would seem like natural methods to use for the analysis of MPRA data, they have not been used for differential analysis of MPRA activity measures. There has been some use of DESeq2 to identify (filter) elements with regulatory activity (differential expression of RNA relative to DNA) (Tewhey et al., 2016; Gisselbrecht et al., 2013). However, these tools have not been used for comparisons of activity measures between groups. In this work we propose the use of log-DNA offsets as potential sensible uses of these software for differential analysis. In our evaluations, we see that this approach is most competitive with mpralm. For the allelic studies (Tewhey et al., 2016; Ulirsch et al., 2016), we observe that the degree of within-sample correlation affects the power of mpralm relative to comparison methods. In particular, there is little difference in the performance of the different methods for the large pool experiment of Tewhey et al. (2016), and this experiment had overall low within-sample correlation. Both the targeted pool experiment of Tewhey et al. (2016) and the Ulirsch experiment had larger within-sample correlations, and we observe that mpralm has increased power over the comparison methods for these datasets. We expect that mpralm will generally be more powerful for paired designs with high within-pair correlations.

In terms of element rankings, mpralm, edgeR, and DESeq2 are similar. However, we observe a substantial difference in ranking between t-tests and mpralm and believe top ranked mpralm elements exhibit better properties compared to those from t-tests.

Linear models come with analytic flexibility that is necessary to handle diverse MPRA designs. Paired designs involving alleles, for example, are easily handled with linear mixed effects models due to computational tractability. The studies we have analyzed here only consider two alleles per locus. It is possible to have more than two alleles at a locus, and such a situation cannot be addressed with paired t-tests, but is easily analyzed using mpralm. This is important because we believe such studies will eventually become routine for understanding results from genome-wide association studies.

While we have focused on characterizing the mpralm linear model framework for differential analysis, it is possible to include variance weights in the multivariate models used in saturation mutagenesis and characterization studies. We expect that modeling the copy number-variance relationship will improve the performance of these models.

For power, we find a substantial impact of even small increases in sample size. This is an important observation because many MPRA studies use 2 or 3 replicates per group, and our results suggest that power can be substantially increased with even a modest increase in sample size. We caution that using less than 4 replicates can be quite underpowered.

In short, the tools and ideas set forth here will aid in making rigorous conclusions from a large variety of future MPRA studies.

### Supplemental Information Availability

The Supplemental Information contains more detailed information on the implementation of mpralm, the running of comparison methods, data processing, and theoretical calculations for the bias and variance of activity estimators.

## Methods

### Count preprocessing

DNA and RNA counts are scaled to have the same library size before running any methods. We perform minimal filtering on the counts to remove elements from the analysis that have low counts across all samples. Specifically, we require that DNA counts must be at least 10 in all samples to avoid instability of the log-ratio activity measures. We also remove elements in which these log-ratios are identical across all samples.

### Modeling

The square root of the standard deviation of the log-ratios over samples is taken as a function of average log DNA levels over samples, and this relationship is fit with a lowess curve. Predicted variances are inverted to form observation-level precision weights. Log-ratios activity measures and weights are used in the voom analysis pipeline. For the allelic studies, a mixed model is fit for each element using the *duplicateCorrelation* module in the limma Bioconductor package (Smyth, Michaud, and Scott, 2005).

### Permutation tests

We performed null permutation experiments to estimate empirical type I error rates (denoted by *α*) at different nominal levels. Specifically, we created permuted sample groups that each were composed half of group 1 samples and half of group 2 samples. In paired experiments, we maintained the linking between samples but swapped group labels.

### Simulation studies to assess accuracy of permutations for error rate estimation

To model MPRA data, we simulated negative binomial data for both DNA and RNA with a range of means and dispersion parameters, and we fix a proportion p to have differential activity between conditions. We simulated both paired and unpaired data to respectively model allelic and environmental studies.

## Acknowledgements

Funding: Research reported in this publication was supported by the National Cancer Institute and the National Institute of General Medical Sciences of the National Institutes of Health under award numbers U24CA180996 and R01GM121459.

## Disclaimer

The content is solely the responsibility of the authors and does not necessarily represent the official views of the National Institutes of Health.

## Conflict of Interest

None declared.

## Supplementary Information

### Supplementary Methods

#### Data

See Table 1. Dataset labels used in figures are accompanied by short descriptions below.

**Melnikov**: Study of the base-level impact of mutations in two inducible enhancers in humans (Melnikov et al., 2012): a synthetic cAMP-regulated enhancer (CRE) and a virus-inducible interferon-beta enhancer (IFNB). We do not look at the IFNB data because it contains only one sample. We consider 3 datasets:

**Melnikov: CRE, single-hit, induced state:** Synthetic cAMP-regulated enhancer, single-hit scanning, induced state.

**Melnikov: CRE, multi-hit, uninduced state:** Synthetic cAMP-regulated enhancer, multi-hit sampling, uninduced state.

**Melnikov: CRE, multi-hit, induced state:** Synthetic cAMP-regulated enhancer, multi-hit sampling, induced state.

**Kheradpour**: Study of the base-level impact of mutations in various motifs (Kheradpour et al., 2013). Transfection into HepG2 and K562 cells.

**Tewhey:** Study of allelic effects in eQTLs (Tewhey et al., 2016). Transfection into two lymphoblastoid cell lines (NA12878 and NA19239) as well as HepG2. In addition two pools of plasmids are considered: a large screening pool and a smaller, targeted pool, designed based on the results of the large pool. We use data from both the large and the targeted pool in NA12878.

**Inoue: chromosomal vs. episomal:** Comparison of episomal and chromosomally-integrated constructs (Inoue et al., 2017). This study uses a wild-type and mutant integrase to study the activity of a fixed set of putative regulatory elements in an episomal and a chromosomally-integrated setting, respectively.

**Ulirsch:** Study of allelic effects in GWAS to understand red blood cell traits (Ulirsch et al., 2016). Transfection into K562 cells as well as K562 with GATA1 overexpressed. We use the data from K562.

**Shen: mouse retina vs. cortex:** Comparison of cis-regulatory elements in-vivo in mouse retina and cerebral cortex (Shen et al., 2016). Candidate CREs that tile targeted regions are assayed in-vivo in these two mouse tissues with adeno-associated virus delivery.

#### Count preprocessing

We use total count normalization to account for differences in library size for both DNA and RNA. Specifically, each count in a sample is divided by that sample’s library size and scaled so that the library size in all samples is the same. We perform minimal filtering on the counts to remove elements from the analysis that have low counts across all samples. Specifically, we require that DNA counts must be at least 10 in all samples to avoid instability of the log-ratio activity measures. We also remove elements in which these log-ratios are identical across all samples. This is necessary for sensible differential analysis. In practice, log-ratios are only identical across all samples if RNA counts are zero across all samples. Both steps also improve the estimation of the copy number-variance relationship used in subsequent modeling by removing clear outliers.

#### Estimating the copy number-variance relationship

After preprocessing the first step is to estimate the copy number-variance relationship that will allow for the estimation of element-specific reliability weights. These weights are ultimately used in element-specific weighted regressions. The square root of the standard deviation of the log-ratios over samples are taken as a function of average log DNA levels over samples, and this relationship is fit with a lowess curve. Predicted variances are inverted to form observation-level precision weights.

#### Modeling

Once the observation-specific weights are calculated, the log-ratios and weights are used in the voom analysis pipeline. If, as in allele-specific activity studies, the different versions of the elements being compared are correlated due to being measured in the same sample, a mixed model is fit for each element using the `duplicateCorrelation` module in the limma Bioconductor package (Smyth, Michaud, and Scott, 2005).

#### Running mpralm, QuASAR, t-test, Fisher’s exact test

For all methods, DNA and RNA counts were first corrected for library size with total count normalization. For edgeR and DESeq2, DNA counts were included as offset terms on the log scale before standard analysis. For the t-test we computed the aggregate estimator of the log-ratio as the outcome measure. For Fisher’s exact test, we summed DNA and RNA counts in the two conditions to form a 2-by-2 table as input to the procedure. For QuASAR-MPRA, we summed RNA counts in each condition to get one reference condition count and one alternative condition count per element. We also summed DNA counts in all samples and in the reference condition to get one DNA proportion for each element. These were direct inputs to the method.

#### Permutation tests

We performed null permutation experiments to estimate empirical type I error rates (denoted by *α*) at different nominal levels. Specifically, we created permuted sample groups that each were composed half of group 1 samples and half of group 2 samples. For example, in a six versus six comparison, we would select three samples from group 1 and three samples from group 2 to be in the first comparison group. The remaining samples would be in the second comparison group. In this way, we expect no differences in activity measures between the comparison groups. In paired experiments, we maintained the linking between samples but swapped group labels to create null comparisons.

#### Estimation of *π*_0_

The proportion of truly null hypotheses for each dataset was estimated using the “lfdr” method in the `propTrueNull` function within limma (Phipson, 2013). This proportion was estimated for mpralm, t-test, QuASAR, edgeR, and DESeq2, and the median of these estimates was used as the estimate for *π*_0_ for that dataset. Fisher’s exact test was excluded from this estimate because it gave an estimate of *π*_0_ that was considerably smaller than the other methods, and which was dubious in light of its uncontrolled type I error rate. These *π*_0_ estimates are used in the FDR calculations of Figure ??.

#### mpralm enables modeling for complex comparisons

While many comparisons of interest in MPRA studies can be posed as a two group comparison (e.g. major allele vs. minor allele), more complicated experimental designs are also of interest. For example, in the allelic study conducted by Ulirsch et al. (2016), putative biallelic enhancer sequences are compared in two cellular contexts. The first is a standard culture of K562 cells, and the second is a K562 culture that induces over-expression of GATA1 for a more terminally-differentiated phenotype. A straightforward question is whether an allele’s effect on enhancer activity differs between cellular contexts. Let *y_eia_* be the enhancer activity measure (log ratio of RNA over DNA counts) for element *e*, in sample *i* for allele a. Let *x*_1*eia*_ be a binary indicator of the mutant allele. Let *x*_2*eia*_ be a binary indicator of the GATA1 over-expression condition. Then the following model

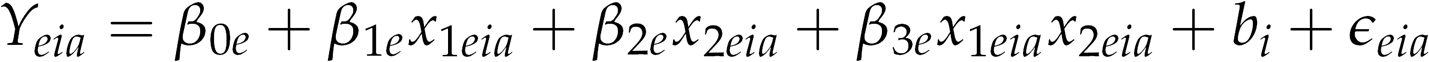

is a linear mixed effects model for activity measures, where *b_i_* is a random effect that induces correlation between the two alleles measured within the same sample. We can perform inference on the *β_3e_* parameters to determine differential allelic effects. Such a model is easy to fit within the mpralm framework, since our framework supports model specifications by general design matrices. In contrast, this question cannot be formulated in the QuASAR, t-test, and Fisher’s exact test frameworks. Neither edgeR nor DESeq2 support the fitting of mixed effects models.

#### Bias and variance of estimators

We use Taylor series arguments to approximate the bias and variance of the aggregate and average estimators. The following summarizes our parametric assumptions:

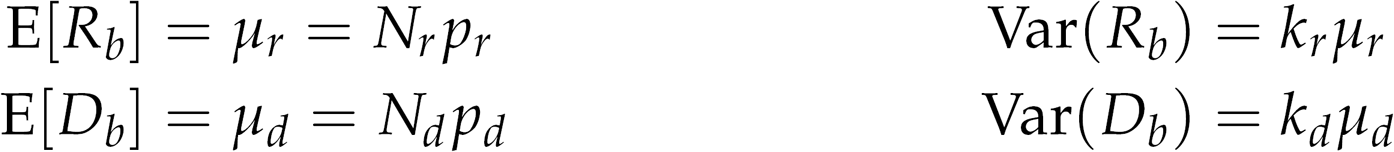

We suppress the dependency of these parameters on sample and element. Library sizes are given by *N*. The fraction of reads coming from a given element is given by *p.* Dispersion parameters are given by *k*. The common library size resulting from total count normalization is given by *L*. The true activity measure of a given element is given by *a* := log(*p_r_*/*p_d_*).

#### Average estimator

The “average estimator” of *a* is an average of barcode-specific log activity measures and is written as:

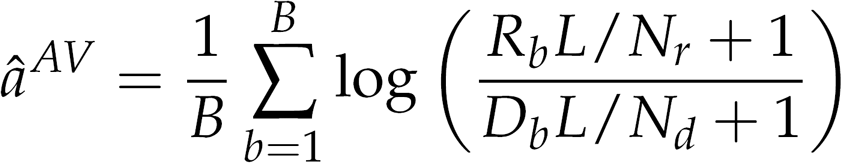

The second-order Taylor expansion of the function

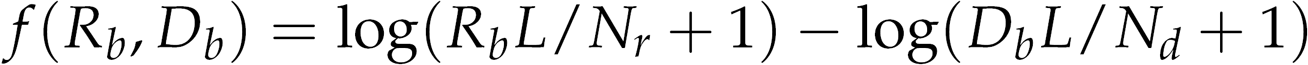

about the point (E[*R_b_*], E[*D_b_*]) = (*μ_r_*, *μ_d_*) is:

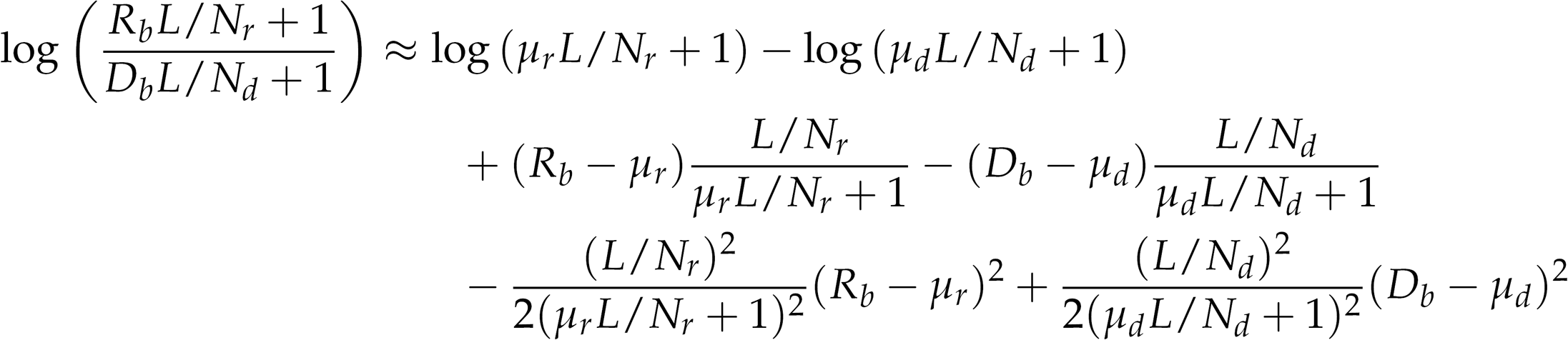

We use the expansion above to approximate the expectation of the average estimator:

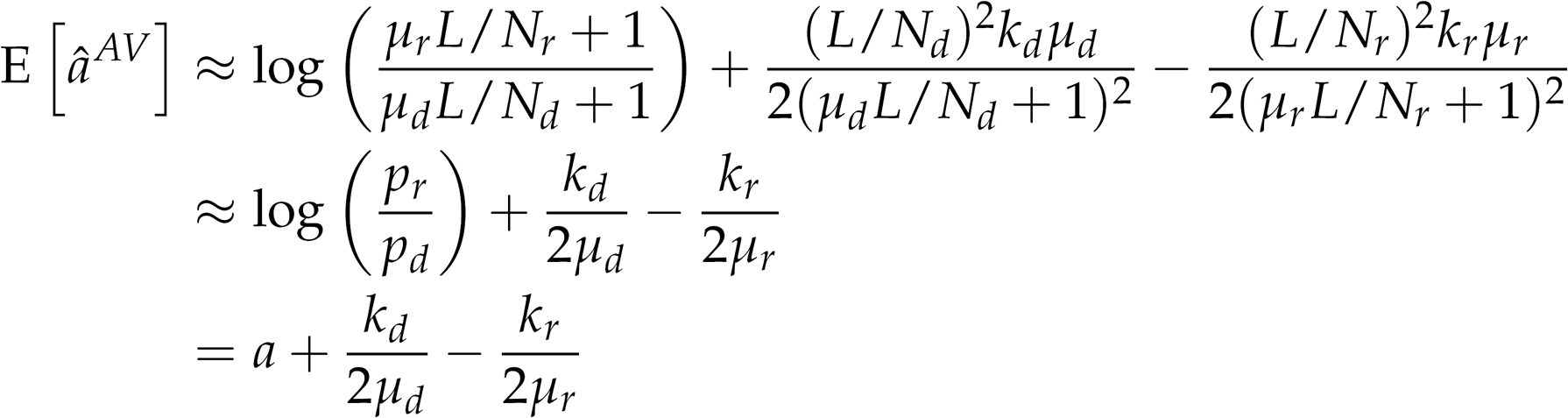

We can also approximate the variance under the assumption that the barcode-specific log-ratios are uncorrelated:

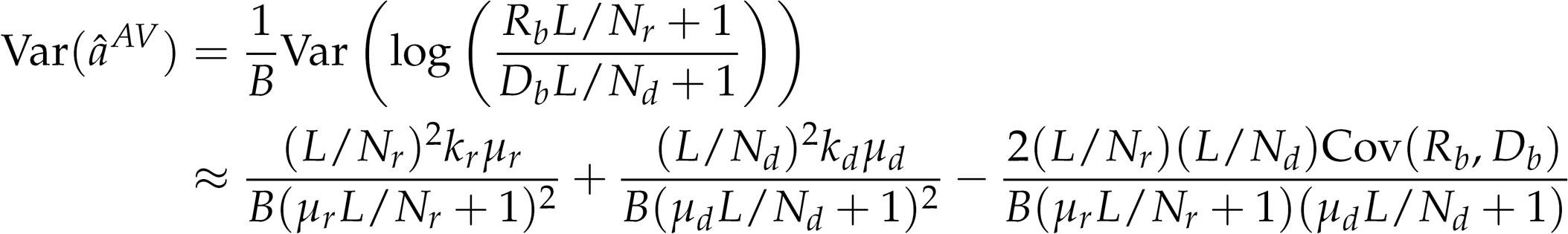

#### Aggregate estimator

The “aggregate estimator” of a first aggregates counts over barcodes and is written as:

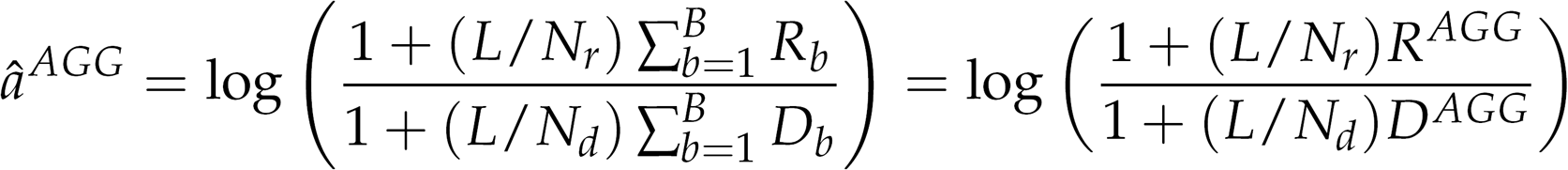

The second-order Taylor expansion of the function

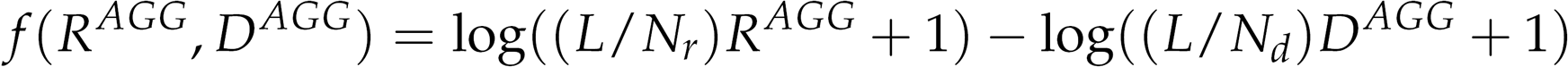

about the point (E[*R^AGG^*],E[*D^AGG^*]) = (*Bμ_r_, Bμ_d_*) is:

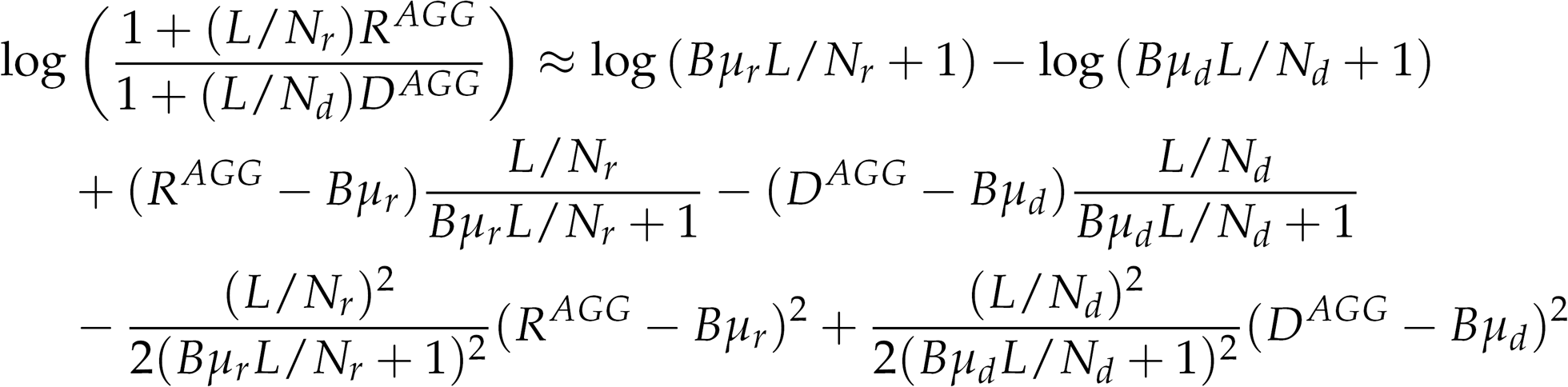

We use the expansion above to approximate the expectation:

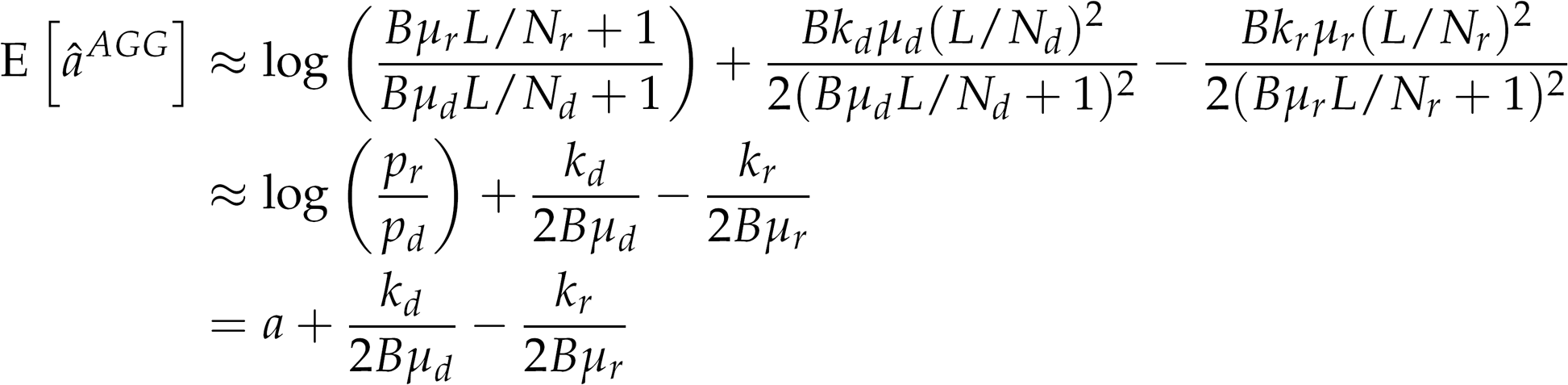

We can also approximate the variance:

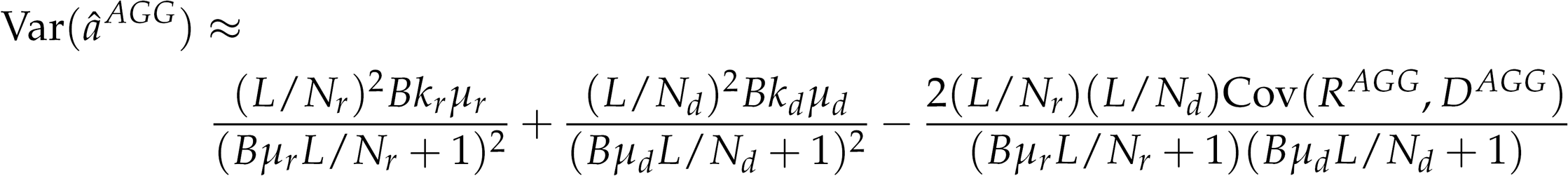

